# Counting cytoplasmic incompatibility factor mRNA using digital droplet PCR

**DOI:** 10.1101/2025.07.30.667682

**Authors:** Lore Van Vlaenderen, William R. Conner, J. Dylan Shropshire

**Author notes:** **Corresponding author** J. Dylan Shropshire. **Email**, (LVV), (WRC), (JDS).

## Abstract

*Wolbachia* bacteria inhabit over half of all insect species and often spread through host populations via efficient maternal transmission and cytoplasmic incompatibility (CI), killing aposymbiotic embryos when fertilized by symbiotic males. *Wolbachia*’s *cifB* gene triggers CI in males, while *cifA*, expressed in females, rescues embryos from CI-induced lethality. In some systems, *cifA* also contributes to CI induction. CI strength—the percentage of embryos that die from CI—is a key determinant of *Wolbachia*’s prevalence in host populations, and *cifB* mRNA levels in testes generally correlate with CI strength. Yet, *cifB*’s rarity can hamper precise quantification, necessitating tissue pooling for reverse transcription quantitative PCR (RT-qPCR) to achieve reliable measurements, obscuring variation at the level of individual insect tissues. Here, we present four RT digital droplet PCR (RT-ddPCR) assays to count rare *cifA* and *cifB* mRNA from *w*Mel *Wolbachia* in *Drosophila melanogaster*. These assays count *cif* transcripts alongside a synthetic spike-in RNA or a *D. melanogaster* housekeeping gene to normalize for technical or biological variation. These assays have a limit of detection of about 1 *cifA* and 3 *cifB* copies per reaction. We expect these methods to be useful for mosquito-control programs that use *w*Mel to block the spread of pathogens from *Aedes aegypti* to humans. Moreover, the oligos were designed with homology to *cifA* and *cifB* sequences from at least 33 *Wolbachia* strains, suggesting utility beyond *w*Mel. These methods will allow researchers to measure *cif* mRNA levels from individual insect tissues, enabling efforts to pair molecular and phenotypic data at unprecedented resolutions.

**Importance:** *Wolbachia*, a maternally transmitted bacterium, is found in over half of all insect species. Its ability to induce cytoplasmic incompatibility (CI), which prevents *Wolbachia*-free eggs from hatching, significantly contributes to its high prevalence in host populations. Public health experts use CI to spread pathogen-blocking *Wolbachia* through mosquito populations, thereby controlling pathogen spread. CI is often weak, resulting in few egg deaths and consequently slowing *Wolbachia*’s spread. We recently discovered that weak CI often correlates with low CI factor B (*cifB*) mRNA levels. However, our understanding of CI-strength variation remains limited because *cifB* is transcribed at low levels, making it challenging to measure in individual insects. Here, we report four RT-ddPCR assays to overcome this challenge. These assays offer high sensitivity for rare targets and maintain accuracy and precision across a wide dynamic range. We expect these tools will enhance efforts to understand CI-strength variation in both natural and applied populations.

## Introduction

Many insects host intracellular bacteria that mothers pass to their offspring (McCutcheon et al., 2019; Moran et al., 2008). Among these microbes, the Alphaproteobacterium *Wolbachia* (Kaur et al., 2021) is one of the most common, found in over half of all insect species (Weinert et al., 2015) and often widespread within host populations (Kriesner et al., 2016). *Wolbachia*’s high prevalence within host populations largely stems from its ability to cause cytoplasmic incompatibility (CI) (Hoffmann et al., 1990; Shropshire et al., 2020). CI occurs when a symbiotic male mates with an aposymbiotic female, killing the resulting embryos (Yen and Barr, 1973). Conversely, embryos from symbiotic females resist CI, giving *Wolbachia*-bearing offspring a selective advantage. Two *Wolbachia* genes orchestrate CI: *cifB*, expressed in testes, induces CI, while *cifA*, expressed in ovaries, rescues embryos from CI-induced lethality (Adams et al., 2021; Cooper et al., 2017; LePage et al., 2017; Shropshire et al., 2018, 2021b; Shropshire and Bordenstein, 2019; Sun et al., 2022). In some systems, both *cifA* and *cifB* must be co-expressed in males to induce CI (e.g., LePage et al., 2017). Beyond its ecological role, public health experts use CI to spread pathogen-blocking *Wolbachia* strains through *Aedes aegypti* mosquito populations, thereby protecting humans from diseases like dengue and Zika (Hoffmann et al., 2024; Simmons et al., 2024; Utarini et al., 2021; Velez et al., 2023).

Strong CI is characterized by high embryo mortality and is crucial for *Wolbachia*’s spread to high frequencies in both natural and applied populations (Hoffmann et al., 1990, 2011; Walker et al., 2011). However, CI strength can vary significantly with host and *Wolbachia* genetics (Cooper et al., 2017; Hughes and Rasgon, 2014; Poinsot et al., 1998; Shropshire et al., 2022; Walker et al., 2011), male age (Awrahman et al., 2014; Reynolds and Hoffmann, 2002; Shropshire et al., 2021a), mating frequency (Awrahman et al., 2014; de Crespigny and Wedell, 2006), diet (Clancy and Hoffmann, 1998; Sinkins et al., 1995), temperature (Bordenstein and Bordenstein, 2011; Ross et al., 2020, 2019), and other factors (reviewed in Shropshire et al., 2020). While a comprehensive understanding of CI-strength variation remains elusive, *cifB*-mRNA levels show a strong, though imperfect, correlation with CI strength across diverse *Wolbachia*-*Drosophila* associations and with male-age-dependent CI in *w*Ri-bearing *D. simulans* (Shropshire et al., 2022, 2021a).

Reverse transcriptase quantitative PCR (RT-qPCR) is traditionally used to measure *cifA* and *cifB* mRNA levels (Shropshire et al., 2022, 2021a). This method requires purifying total RNA, synthesizing complementary DNA (cDNA) through reverse transcription, and then performing PCR with the inclusion of a fluorescent DNA-binding probe or an intercalating dye like SYBR Green to monitor amplicon accumulation. Each PCR cycle doubles the target amplicon abundance, which in turn doubles the fluorescent signal. Researchers infer the initial quantity of target cDNA by identifying the cycle at which its fluorescent signal crosses a predetermined threshold. While *cifA* mRNA is typically abundant and readily measured by RT-qPCR, *cifB* transcripts can be rare (Gutzwiller et al., 2015; Lindsey et al., 2018). Detecting *cifB* commonly requires more than 30 amplification cycles to reach the detection threshold, even when cDNA is derived from samples containing tissues pooled from 15 or more individuals (Shropshire et al., 2022, 2021a). These expression dynamics hinder efforts to correlate molecular data with phenotypic outcomes in individual insects and complicate studies in systems where *cifB* transcripts are especially rare.

Compared to RT-qPCR, reverse transcriptase digital droplet PCR (RT-ddPCR) offers enhanced sensitivity, accuracy, and precision (Vogelstein and Kinzler, 1999). Initially, RT-ddPCR assays share steps with RT-qPCR, including RNA extraction and purification, cDNA synthesis, and setting up PCR reactions with a fluorescent DNA-binding probe or an intercalating dye. However, RT-ddPCR differs significantly in several key ways. First, the reaction is partitioned into thousands of nanoliter-sized droplets. Second, PCR is performed on each droplet for the maximum intended number of cycles. Finally, a microfluidic device with a fluorescence detector measures the fluorescence of each droplet, classifying droplets as positive or negative for the target based on high or low fluorescence intensity, respectively. Assuming targets are evenly distributed across droplets, and that some droplets lack target molecules, Poisson statistics can be applied to estimate the number of targets in each positive droplet. Therefore, RT-ddPCR directly counts the number of target molecules, rather than inferring abundance from variation in fluorescence levels across amplification cycles.

RT-ddPCR’s benefits stem from its endpoint detection and partitioning capabilities. Unlike RT-qPCR, which assumes 100% reaction efficiency and infers cDNA abundance from exponential doubling, RT-ddPCR simply measures the presence or absence of a target once the reaction is complete. This eliminates the unrealistic assumption of perfect efficiency, making RT-ddPCR more resistant to PCR inhibitors often introduced during RNA purification and cDNA synthesis. Consequently, this enhances the assay’s accuracy and precision. Moreover, RT-ddPCR significantly boosts measurement confidence. While it is common in RT-qPCR to perform two or three technical replicates per sample, each of the thousands of nanoliter-sized droplets in an RT-ddPCR reaction serves as an individual replicate. This vastly increases the number of technical replicates, leading to greater confidence in the results. Since the limit of detection is determined by the upper limit of the 95% confidence interval from negative controls, this improved confidence directly enhances assay sensitivity. For example, assuming no positive droplets in the negative control, the limit of detection for an RT-ddPCR assay is three target copies from 3,000 droplets with 95% confidence (Hindson et al., 2011; Hoshino and Inagaki, 2012; Pinheiro et al., 2012). The QX200 ddPCR system used in this study reliably generates between 10 and 20 thousand droplets per reaction, further bolstering confidence.

Here, we present optimized methods for the extraction, purification, and processing of RNA from individual *D. melanogaster* testes to count *cifA* and *cifB* mRNA of *w*Mel *Wolbachia* using two-step RT-ddPCR. While ddPCR has been used to measure *Wolbachia* abundance in multiple studies (Fisher et al., 2019; Kakumanu et al., 2024, 2024; Kilpatrick et al., 2024; Njogu et al., 2025; Njogu and Shropshire, 2024), to our knowledge, this is the first application of RT-ddPCR to count *Wolbachia* mRNA. For each *cif* gene, we developed two duplex RT-ddPCR assays: *cif*/spike and *cif*/*β-Spec*. The *cif*/spike assays simultaneously measure *cif* mRNA levels and RNA processing efficiency using a spike-in RNA, enabling users to account for technical variation between samples. These *cif*/spike assays have a limit of detection of 1 *cifA* and 3 *cifB* copies per 20 µL reaction. In contrast, the *cif*/*β-Spec* assays normalize *cif* transcription against that of *D. melanogaster*’s housekeeping gene *β-Spectrin* (FlyBase ID FBgn0250788) to control for biological variation in transcription, with a limit of detection of 1 *cifA* and 1 *cifB* copy per reaction. We expect the *cif*/spike assays to offer broad utility beyond *w*Mel-bearing *D. melanogaster*. The *w*Mel *Wolbachia* strain is used in *Ae. aegypti* for mosquito-borne disease control, and we designed the *cif* RT-ddPCR oligos with homology to 39 *cifA* (from 32 *Wolbachia* strains) and 34 *cifB* variants (from 27 *Wolbachia* strains). These methods will significantly enhance the ability to measure *cif* mRNA in individual insects, particularly when *cif* genes are expressed at low levels.

## Results

### Optimizing RNA purification for low-biomass samples

To enable sensitive RT-ddPCR assays for rare *cifA* and *cifB* mRNA in individual insects, we first developed and validated an RNA extraction protocol optimized for low-biomass samples. Due to its documented high RNA yields, we decided to extract and purify RNA through phase separation (Tesfamichael et al., 2020; Zhao et al., 2023). In brief, the protocol (detailed in Van Vlaenderen and Shropshire, 2025) involves bead-mill homogenization, TRIzol:chloroform phase separation, and isopropanol-ethanol RNA precipitation with a glycogen carrier. A final ethanol reprecipitation step significantly improves RNA purity by reducing both protein (average A260/A280 improved from 1.87 to 1.94; Paired *T*-test *P* = 0.041; *N* = 9) and chemical contamination (average A260/A230 improved from 0.93 to 1.45; Paired *T*-test *P* = 9.8e-3; *N* = 9) without significantly impacting RNA yield (Paired *T*-test *P* = 0.44; *N* = 9) (**Data S1**).

We tested this protocol’s accuracy and precision across samples with different biomass by extracting RNA from 1, 4, 10, and 20 pairs of *w*Mel-bearing *D. melanogaster* testes. RNA yield per pair of testes is consistent regardless of the number of testes in the sample (*R*^*2*^ = 0.069, *P* = 0.5), averaging 2.1 ng/μL per pair of testes in 25 μL of low EDTA buffer (52.5 ng total; **Fig 1A**). Consequently, the total RNA concentration significantly increases with the number of testes (Pearson’s *R*^*2*^ = 0.86, *P* = 2.9e-4; **Fig 1B**). The observed slope of this relationship is consistent with the average yield per testis pair (One sample *T*-test *P* = 0.79), supporting accurate recovery across biomass levels. However, sample-to-sample variation in RNA yield is higher in groups with a higher number of testes per sample (Pearson’s *R*^*2*^ = 0.99, *P* = 4.8e-3; **Fig 1C**), indicating reduced precision as biomass increases. These findings demonstrate that this RNA extraction and purification protocol yields about 52.5 ng of RNA per pair of testes across a range of tissue abundances, and is especially precise when extracting from low-biomass samples.

**Figure 1.**
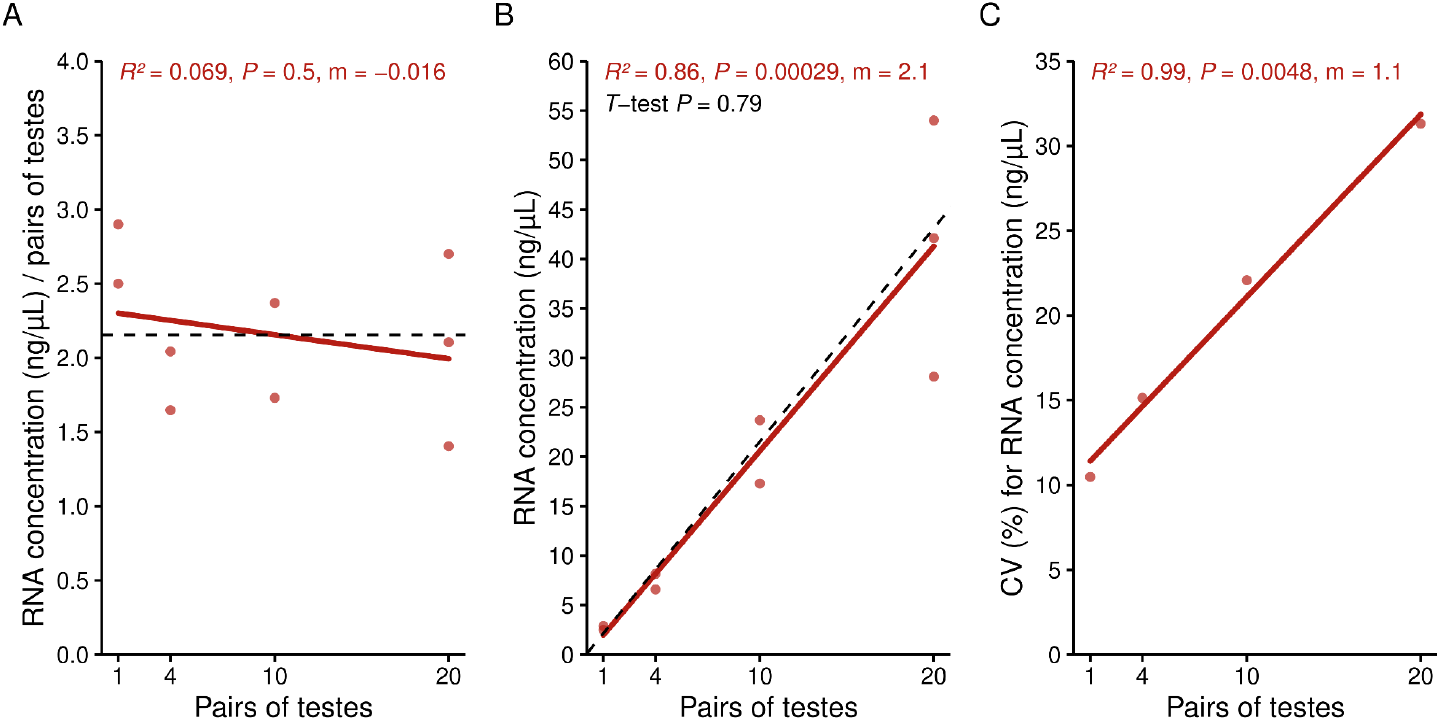
Phase-separation RNA extraction consistently yields about 2.1 ng/µL of RNA per pair of testes. **(A)** RNA concentration per pair of testes in 25 μL of low EDTA buffer remains consistent regardless of the number of testes in the sample. The dashed black line represents the average RNA yield per pair of testes across all extractions, 2.1 ng/µL. **(B)** Total RNA concentration significantly increases with the number of testes per sample. The dashed black line indicates the expected linear relationship between total RNA yield and the number of testes, calculated using the average RNA yield per pair of testes as the slope. We used a one-sample *T*-test to compare the observed and expected slopes. **(C)** Variability in sample-to-sample RNA yield, as measured by the coefficient of variation (CV), is significantly higher in sample groups containing more testes. **(A-C)** We quantified RNA concentrations using a Qubit 4 Fluorometer with the Qubit RNA High Sensitivity Kit. Each plot displays the Pearson’s correlation coefficient, the corresponding *P*-value, and the regression line’s slope (m). Raw data for these analyses are available in **Data S1**.

### Developing generalizable *cif* RT-ddPCR assays

To design primers and probes for the *cifA* and *cifB* mRNA of the *w*Mel *Wolbachia* strain from *D. melanogaster*, we first gathered a large set of *cifA* and *cifB* sequences using BLAST against an in-house *Wolbachia* genome database. Next, we constructed multiple sequence alignments (MSAs), generated consensus sequences with variable nucleotide masks, and used Primer3Web to design forward/reverse primers and a fluorescein amidite (FAM)-labeled probe. We iteratively refined the MSAs, selecting sequences based on their similarity to *w*Mel’s *cifA* and *cifB* genes until we identified optimal oligos (see Materials & Methods). Finally, we validated the primers against the NCBI nr database using NCBI Primer-BLAST. This confirmed no off-target binding to humans (taxid: 9606), *Drosophila* (taxid: 7215), and *Wolbachia* (taxid: 953).

We also used Primer-BLAST to identify homology across various *Wolbachia* genomes. The *cifA* oligos show homology to 39 sequences (**Data S2**), and the *cifB* oligos to 34 sequences (**Data S3**), spanning 30 Supergroup A and 3 Supergroup B *Wolbachia* strains. This broad homology includes *Wolbachia* in 12 Diptera from the family Drosophilidae: *D. ananassae* (*w*Ana), *D. biauraria* (*w*Biau), *D. chauvacae* (*w*Ack), *D. incompta* (*w*Inc), *D. innubila* (*w*Inn), *D. melanogaster* (*w*Mel), *D. santomea* (*w*San), *D. simulans* (*w*Ri, *w*Ha), *D. sturtevanti* (*w*Stv), *D. teissieri* (*w*Tei), *D. yakuba* (*w*Yak), and *Zaprionus tsacasi* (*w*Zts). Beyond the Drosophilidae, the oligos were homologous to *cif* genes from *Wolbachia* in 5 Syrphidae flies (*Brachyopa scutellaris, Cheilosia soror, Epistrophe grossularia, Microdon myrmicae*, and *Volucella bombylans*), 3 Tachinidae flies (*Lypha dubia, Rhagoletis cingulata* (*w*Cin), and *Sturmia bella*), 2 Muscidae flies (*Haematobia irritans* (*w*Irr) and *Limnophora tigrina*), 5 other Diptera (*Coremacera marginata, Delia radicum, Liromyza huidobrensis* (wLtri), *Opomyza germinationis*, and *Protocalliphoroa azurea*), 3 Hymenoptera (*Aporus unicolor, Ectemnius continuus*, and *Ophion costatus*) and 3 Lepidoptera (*Hofmannophila pseudospretella, Pheosia gnoma*, and *Rhopobota naevana*). Notably, *cifA* oligos are not homologous to *cifs* in *Cheilosia soror*. Similarly, *cifB* oligos lack homology to *cifs* in *D. biauraria, Ectemnius continuus, Epistrophe grossularia, Limnophora tigrina, Microdon myrmicae*, and *Rhopobota naevana*.

### Evaluating RT-ddPCR droplet differentiation

We developed duplex RT-ddPCR assays using the newly designed *cif* oligos and a commercially available RNA spike-in control kit with hexachlorofluorescein (HEX)-labelled probes (TATAA Biocenter, RS25SI). These assays simultaneously count *cifA* or *cifB* and a synthetic spike-in RNA molecule. For both the *cifA*/spike and *cifB*/spike assays, droplets separate into two clear populations on the FAM channel, indicating the presence or absence of *cifA* and *cifB* (**Fig 2A, B**). However, the HEX channel, which measures the spike-in sequence, exhibits three droplet clusters for both assays. In the *cifA*/spike assay, an intermediate HEX cluster appears close to the positive spike-in cluster (**Fig 2C**). Conversely, in the *cifB*/spike assay, the intermediate HEX cluster is positioned near the negative spike-in cluster (**Fig 2D**). Simultaneously analyzing FAM and HEX signals offer crucial insights into these intermediate clusters (**Fig 3A**). For the *cifA*/spike assay, droplets in the upper HEX cluster have a low FAM signal, while droplets in the intermediate cluster have a high FAM signal (**Fig 3A**). This indicates that the intermediate HEX cluster in the *cifA*/spike assay contains both the spike-in control and *cifA*. In contrast, for the *cifB*/spike assay, the intermediate HEX cluster corresponds exclusively to droplets positive for only *cifB* (**Fig 3B**). Therefore, these intermediate droplets in the *cifB*/spike assay are negative for the spike-in sequence. These findings demonstrate that we can effectively differentiate droplets containing no template, a single template (*cif* or spike-in), or both templates based on their distinct fluorescence profiles.

**Figure 2.**
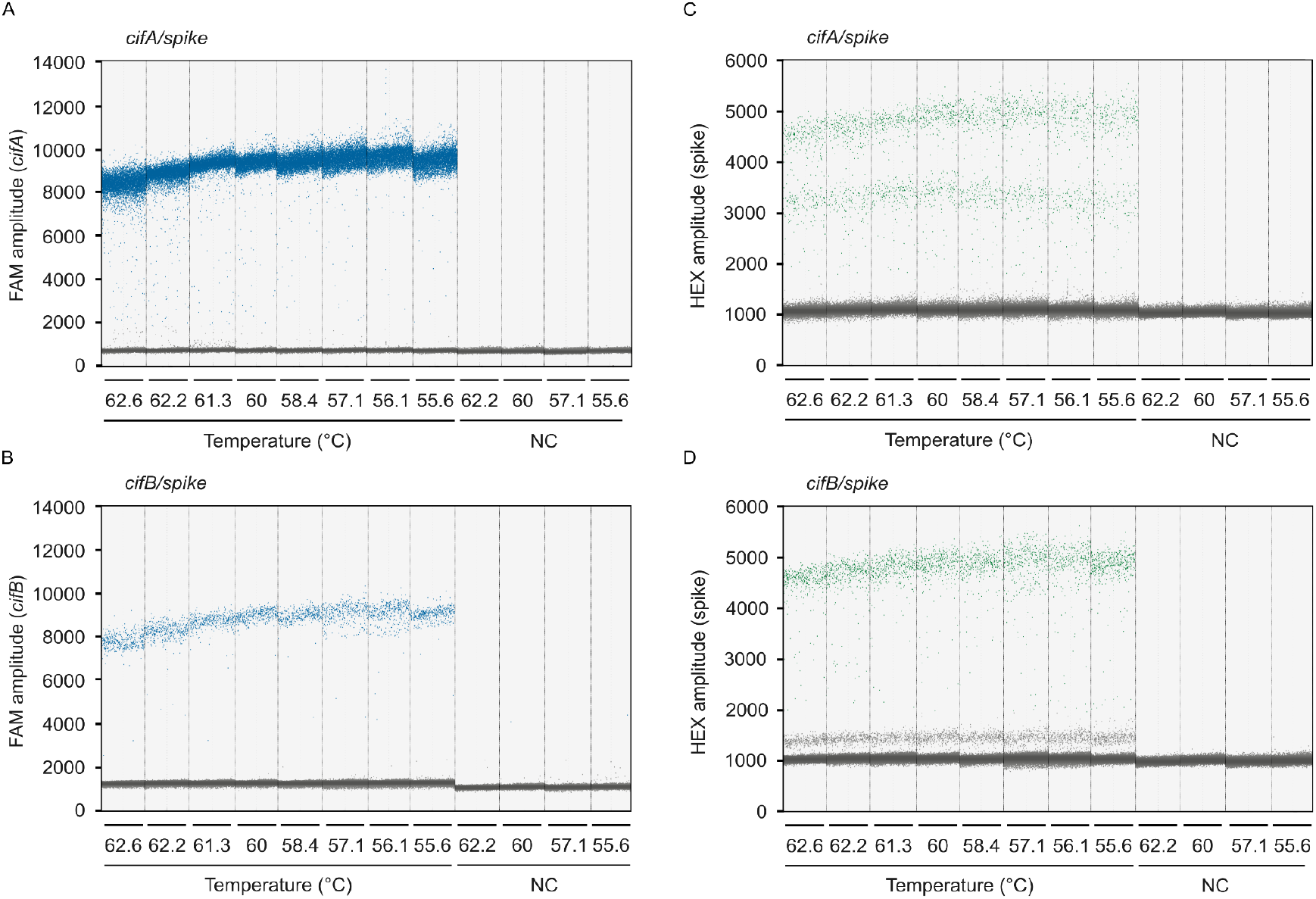
*cif*/spike RT-ddPCR droplet discrimination is robust across annealing temperatures. **(A)** *cifA* detection on the FAM channel is consistent across a range of annealing temperatures (62.6°C to 55.6°C) in the *cifA*/spike assay. **(B)** *cifB* detection on the FAM channel is consistent across the same annealing temperatures in the *cifB*/spike assay. Detection for a synthetic spike-in sequence is consistent across annealing temperatures in both the **(C)** *cifA*/spike and **(D)** *cifB*/spike assays. Vertical dotted lines delineate the results from 20 μL ddPCR reactions, each containing 2 μL of cDNA template derived from RNA purified from 20 pairs of testes. Data points represent individual droplets, with each reaction analyzed containing no fewer than 10,000 droplets. Droplets exhibiting high amplitude (blue on FAM; green on HEX) indicate the presence of the target template, while low amplitude droplets (gray) represent the absence of the template. NC; no-template control. Raw data are available in **Data S4** and **Data S5**.

**Figure 3.**
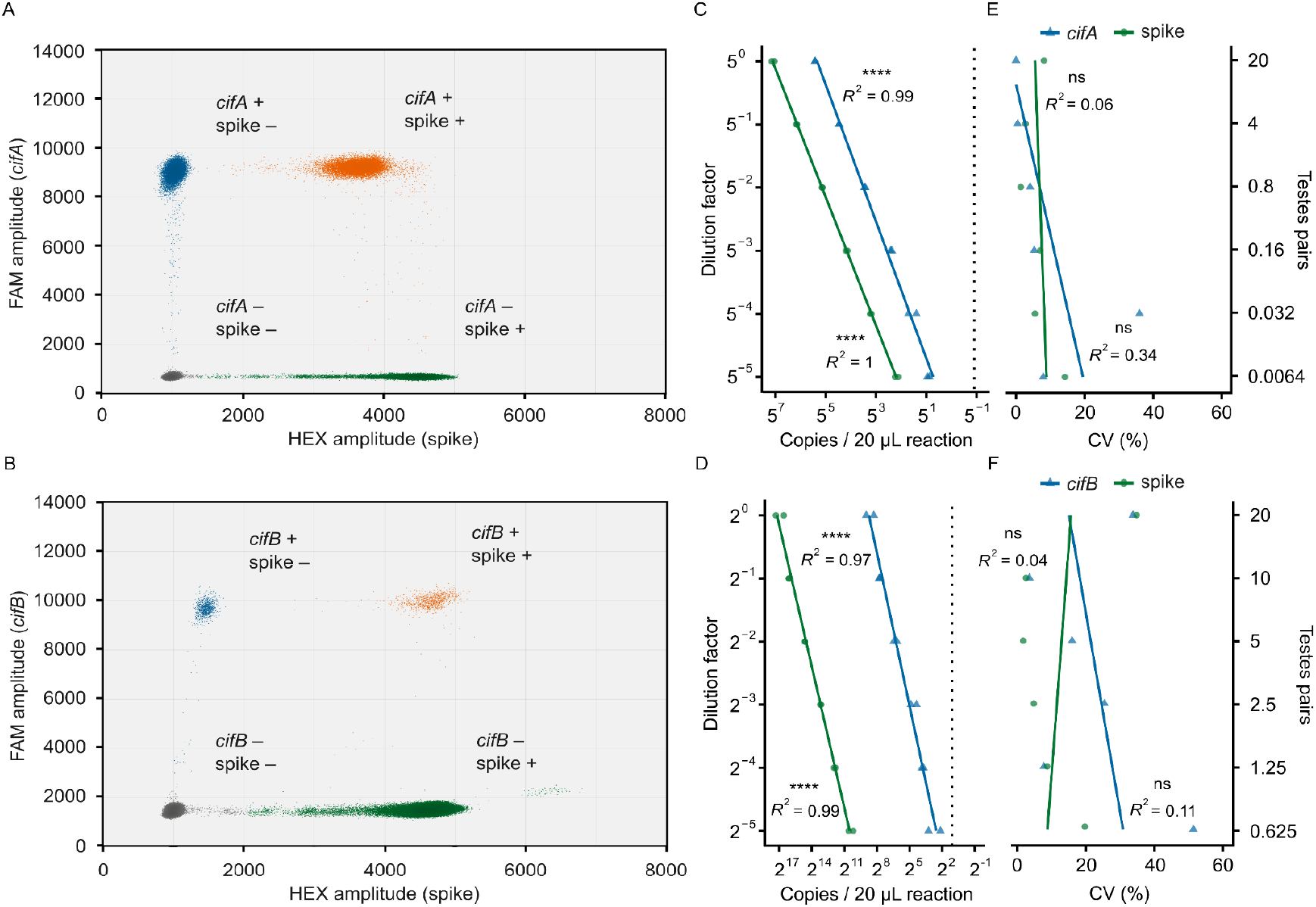
The *cif*/spike RT-ddPCR assays are accurate and precise. **(A, B)** Droplets from 12 combined **(A)** *cifA*/spike and **(B)** *cifB*/spike RT-ddPCR reactions reveal that droplets cluster into four distinct groups based on FAM and HEX amplitudes. Droplets are colored to indicate their content: gray (no targets), blue (*cif* only), green (spike only), and orange (both targets). We calculated target concentrations from both single- and double-positive droplets. **(C, D)** A strong linear relationship between the dilution factor and detected abundance of both targets in **(C)** *cifA*/spike and **(D)** *cifB*/spike RT-ddPCR reactions confirms both assays are accurate across a broad dynamic range. Each 20 µL RT-ddPCR reaction contained 2 µL cDNA and yielded no fewer than 10,000 droplets. Vertical dotted lines indicate the *cif* limits of detection, based on the upper limit of the 95% confidence interval from the no-template controls. **(E, F)** Coefficients of variation (CV) do not significantly vary with concentration in the **(E)** *cifA*/spike or **(F)** *cifB*/spike assays. **(C–F)** Statistical significance is denoted as: *P* > 0.05 (ns), *P* ≤ 0.05 (*), *P* ≤ 0.01 (**), *P* ≤ 0.001 (***), *P* ≤ 0.0001 (****). Statistical tests are Pearson’s product–moment correlations. Raw data are available in **Data S6** and **Data S7**.

### Testing RT-ddPCR efficiency across annealing temperatures

To determine the optimal annealing temperature for the *cif*/spike RT-ddPCR assays, we tested RT-ddPCR performance across a temperature gradient ranging from 56.2°C to 62.2°C. The optimal temperature was defined as that which produced the largest amplitude difference between positive and negative droplets. Fluorescence amplitudes of negative droplets are consistent across temperature treatments for *cifA* (FAM; 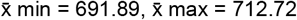 ; **Fig 2A**), *cifB* (FAM; 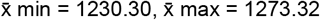; **Fig 2B**), and spike (HEX; 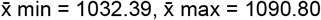; **Fig 2C,D**). Conversely, positive droplet signal amplitude depends on temperature; we detect the highest amplitude for *cifA* at 56.1°C (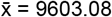; **Fig 2A**), *cifB* at 56.1°C (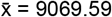; **Fig 2B**), and spike at 55.6°C (*cifA*/spike; 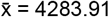 **Fig 2C**) and 58.4°C (*cifB*/spike; 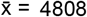; **Fig D**). However, the separation between positive and negative droplet clusters is unambiguous at all tested temperatures. Based on this performance, we selected 60°C as the standard annealing temperature for all subsequent experiments.

### Validating *cif*/spike RT-ddPCR assays

To assess the accuracy of the *cif*/spike assays, we tested each assay against a cDNA dilution series. We derived the cDNA from RNA purified from three samples, each containing 20 pairs of *Drosophila* testes. We used a 1:5 dilution factor for the *cifA*/spike assays and a 1:2 factor for the *cifB*/spike assays. We performed RT-ddPCR across these dilutions using 2 µL of cDNA per 20 µL reaction, and calculated target concentrations from both single-positive and double-positive droplets. For these assays to be practical for analyzing single testes pairs, we needed to reliably detect *cifA* at the 5^-2^ dilution (equivalent to 0.8 testes pairs) and *cifB* at the 2^-5^ dilution (equivalent to 0.625 testes pairs). We detect 374.86 *cifA* copies/reaction at the 5^-2^ dilution (**Fig 3C**) and 8.20 *cifB* copies/reaction at the 2^-5^ dilution (**Fig 3D**). These values are above the limit of detection, defined by the upper limit of the 95% confidence interval for the no-template controls (0.54 *cifA* and 2.59 *cifB* copies/reaction, respectively). Furthermore, we observe a strong, positive linear relationship between the dilution level and the measured abundance for both the *cifA*/spike assay (*cifA* Pearson’s *R*^*2*^ = 0.99, *P* = 6.88e-12; spike Pearson’s *R*^*2*^ = 1, *P* = 2.21e-17; **Fig 3C**) and the *cifB*/spike assay (*cifB* Pearson’s *R*^*2*^ = 0.97, *P* = 5.16e-9; spike Pearson’s *R*^*2*^ = 0.99, *P* = 3.23e-11; **Fig 3D**). Collectively, these data confirm that both *cifA*/spike and *cifB*/spike assays are suitable for counting mRNA from individual testes pairs and accurately measuring the abundance of their respective targets across a broad range of concentrations.

We evaluated assay precision by calculating the coefficient of variation between technical replicates (*N* = 2) at each dilution factor. As expected (see Discussion), precision declined as target concentrations decreased for *cifA* and *cifB*, although not significantly. For instance, the coefficient of variation for *cifA* in the *cifA*/spike assay increased from 0.03% at the 5^0^ dilution to 7.98% at the 5^-5^ dilution (Pearson’s *R*^*2*^ = 0.338, *P* = 0.226; **Fig 3C**). Similarly, the coefficient of variation for *cifB* in the *cifB*/spike assay rose from 12.29% at the 2^0^ dilution to 33.14% at the 2^-5^ dilution (Pearson’s *R*^*2*^ = 0.109, *P* = 0.522; **Fig 3D**). We observed similar patterns for the spike-in sequence in both assays (*cifA*/spike Pearson’s *R*^*2*^ = 0.0615, *P* = 0.635; *cifB*/spike Pearson’s *R*^*2*^ = 0.0387, *P* = 0.709; **Fig 3C, D)**. These data demonstrate that the *cif*/spike assays exhibit good precision, albeit with lower precision as targets are made rarer.

### Validating *cif*/*β-Spec* RT-ddPCR assays

In addition to the *cif*/spike RT-ddPCR assays, we developed duplex RT-ddPCR assays to simultaneously count *cifA* or *cifB* and a *D. melanogaster* housekeeping gene *β-Spec*. Similar to the *cif*/spike assays, representative reactions for both the *cifA*/*β-Spec* (**Fig 4A**) and *cifB*/*β-Spec* (**Fig 4B**) assays demonstrate clear separation of droplet populations, enabling unambiguous quantification of both targets. We assessed the accuracy of the *cif*/*β-Spec* assays using the same methodology as the *cif*/spike assays. In brief, we prepared serial cDNA dilutions (a 1:5 dilution factor for *cifA*/*β-Spec* and 1:2 for *cifB*/*β-Spec*) and performed RT-ddPCR using 2 µL of cDNA per 20 µL reaction. For these assays to be practical for counting *cif* mRNA from individual testes pairs, they needed to reliably detect *cifA* at the 5^-2^ dilution and *cifB* at the 2^-5^ dilution. We detect 356.79 *cifA* copies/reaction (**Fig 4C**) and 8.13 *cifB* copies/reaction (**Fig 4D**) at these respective concentrations. These values are above the limits of detection, defined by the upper limit of the 95% confidence interval of the no-template controls (0.59 *cifA* and 0.87 *cifB* copies/reaction). Consistent with the *cif*/spike assay results, dilution factor and the measured abundance for both the *cifA*/*β-Spec* assay (*cifA* Pearson’s *R*^*2*^ = 0.985, *P* = 1.94e-10; *β-Spec* Pearson’s *R*^*2*^ = 0.986, *P* = 1.59e-12; **Fig 4C**) and the *cifB*/spike assay (*cifB* Pearson’s *R*^*2*^ = 0.98, *P* = 1.52e-11; *β-Spec* Pearson’s *R*^*2*^ = 0.998, *P* = 2.91e-17; **Fig 4D**) are strongly correlated. Collectively, these data confirm that, similar to the *cif*/spike assays, both the *cifA*/*β-Spec* and *cifB*/*β-Spec* assays are well-suited for detecting mRNA from individual testes pairs and accurately measuring target abundance across a wide dynamic range.

**Figure 4.**
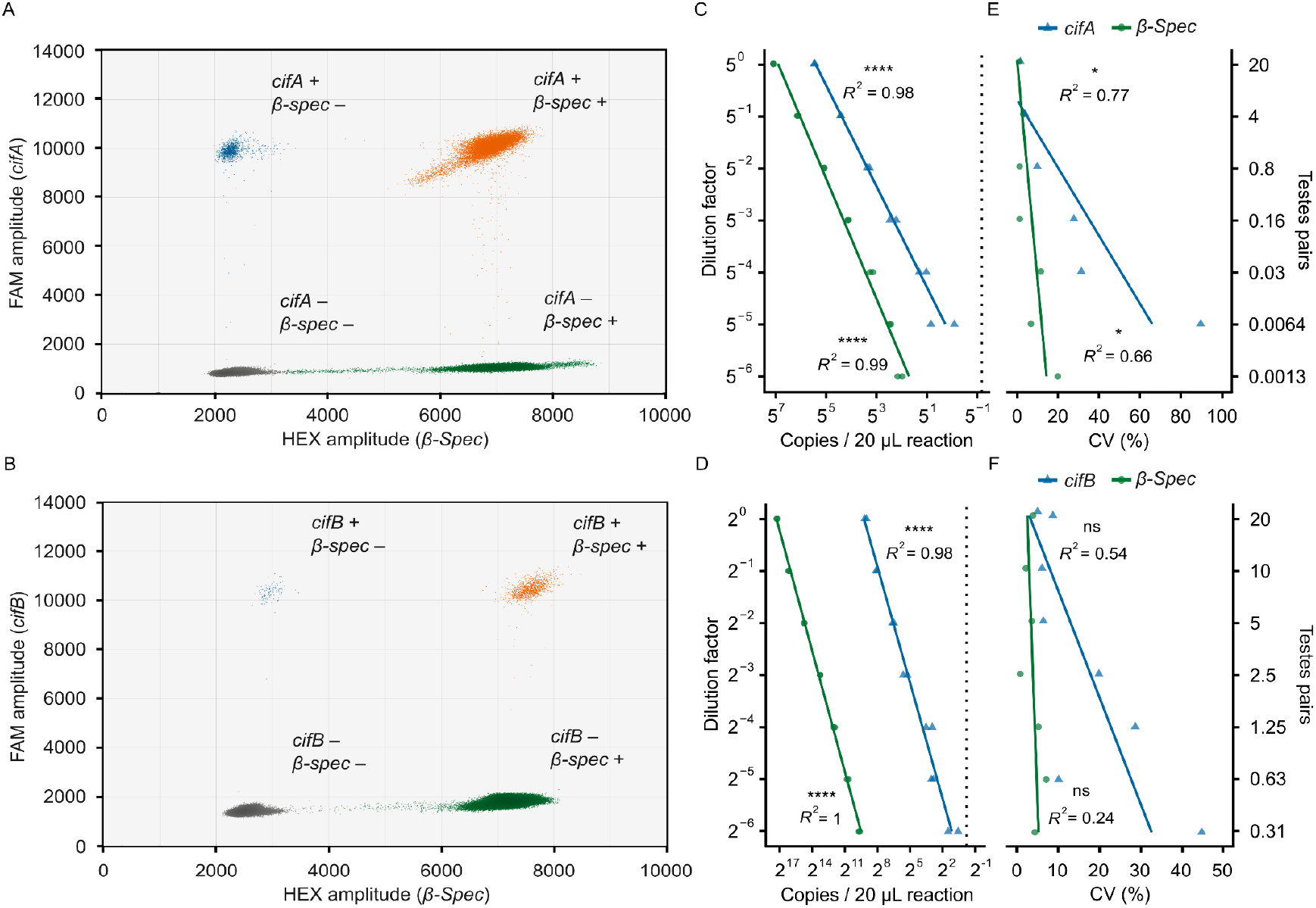
The *cif*/*β-Spec* RT-ddPCR assays are accurate and precise. **(A, B)** Droplets from 14 combined **(A)** *cifA*/*β-Spec* and **(B)** *cifB*/*β-Spec* RT-ddPCR reactions reveal that droplets cluster into four distinct groups based on FAM and HEX amplitudes. Droplets are colored to indicate their content: gray (no targets), blue (*cif* only), green (*β-Spec* only), and orange (both targets). We calculated target concentrations from both single- and double-positive droplets. **(C, D)** A strong linear relationship between the dilution factor and detected abundance of both targets in **(C)** *cifA*/*β-Spec* and **(D)** *cifB*/*β-Spec* RT-ddPCR reactions confirms both assays are accurate across a broad dynamic range. Each 20 µL RT-ddPCR reaction contained 2 µL cDNA and yielded no fewer than 10,000 droplets. Vertical dotted lines indicate the *cif* limits of detection, based on the upper limit of the 95% confidence interval from the no-template controls. **(E, F)** Coefficients of variation (CV) significantly increase as samples are diluted in the **(E)** *cifA*/*β-Spec* assay but not the **(F)** cifB/*β-Spec* assay. **(C–F)** Statistical significance is denoted as: *P* > 0.05 (ns), *P* ≤ 0.05 (*), *P* ≤ 0.01 (**), *P* ≤ 0.001 (***), *P* ≤ 0.0001 (****). Statistical tests are Pearson’s product–moment correlations. Raw data are available in **Data S8** and **Data S9**.

Next, we assessed the precision of the *cif*/*β-Spec* assays by calculating the coefficient of variation between technical replicates (*N* = 2) for each dilution. The coefficients of variation for *cifA* significantly increase from 1.64% at the 5^0^ dilution to 89.74% at the 5^-6^ dilution (Pearson’s *R*^*2*^ = 0.771, *P* = 0.0215; **Fig 4E**). Similarly, the coefficients of variation for *cifB* increase, though not significantly, from 8.01% at the 2^0^ dilution to 43.57% at the 2^-5^ dilution (Pearson’s *R*^*2*^ = 0.54, *P* = 0.0584; *β-Spec* Pearson’s *R*^*2*^ = 0.24, *P* = 0.266; **Fig 4F**). For *β-Spec*, we observed a significant increase in coefficient of variation only in the *cifA*/*β-Spec* assay, which encompassed a broader concentration range (Pearson’s *R*^*2*^ = 0.655, *P* = 0.0274; **Fig 4E**). These data confirm that the *cif*/*β-Spec* assays are precise, with a predictable decline in precision when counting low-abundance targets.

### Evaluating DNase-treatment kits

Since we confirmed the *cif*/spike and *cif*/*β-Spec* assays are accurate and precise even for rare transcripts, minimizing genomic DNA (gDNA) carryover from RNA purifications becomes crucial. To establish a suitable methodology, we first extracted and pooled RNA from ten individual males, creating two large-volume samples. We then evenly divided each pooled sample and treated their aliquots with one of five different protocols from two Invitrogen kits: DNA-free and TURBO DNA-free. Both kits offered “routine” protocols, which involved adding 1 µL of DNase to the sample, incubating at 37°C for 30 minutes, and then inactivating the DNase with an included EDTA-containing reagent. The “rigorous” protocols for both kits required adding an additional 1 µL of DNase and repeating the incubation before inactivation. For the DNA-free kit, we also tested a “rigorous x2” protocol, which performed the rigorous treatment twice.

After DNase-treatment, we assessed gDNA presence using two approaches. First, we amplified a non-transcribed region of *D. melanogaster*’s 28s rDNA using standard PCR and visualized the amplicon via gel electrophoresis. We observe a 28s-associated amplicon when using the DNA-free routine kit; all other protocols yield no detectable DNA (**Fig 5A**). Second, we used the newly designed *cifA*/*β-Spec* RT-ddPCR assay to measure the amount of gDNA contamination. In non-DNase-treated RNA samples, we measure 32,141 *cifA* and 812 *β-Spec* copies per reaction, indicating high levels of gDNA contamination. All five DNase-treatment protocols significantly reduce the concentration of *cifA* by at least 95.15% and *β-Spec* by at least 96.93% (**Fig 5B**). However, gDNA is still detected with the DNA-free routine (1,558 *cifA* and 25 *β-Spec* copies per reaction) and rigorous (548 *cifA* and 6 *β-Spec* copies per reaction) protocols, suggesting incomplete DNA digestion. In contrast, we detect 0 *cifA* copies per reaction with the DNA-free rigorous x2 protocol and both TURBO protocols. Only the TURBO rigorous protocol completely removed *β-Spec* gDNA contamination. Among the three protocols that completely remove *cifA* gDNA, *cifA* and *β-Spec* concentration are similar after reverse transcription (**Fig 5C**). Therefore, we conclude that treating purified RNA from testes extracts with the TURBO rigorous protocol is sufficient to digest all gDNA.

**Figure 5.**
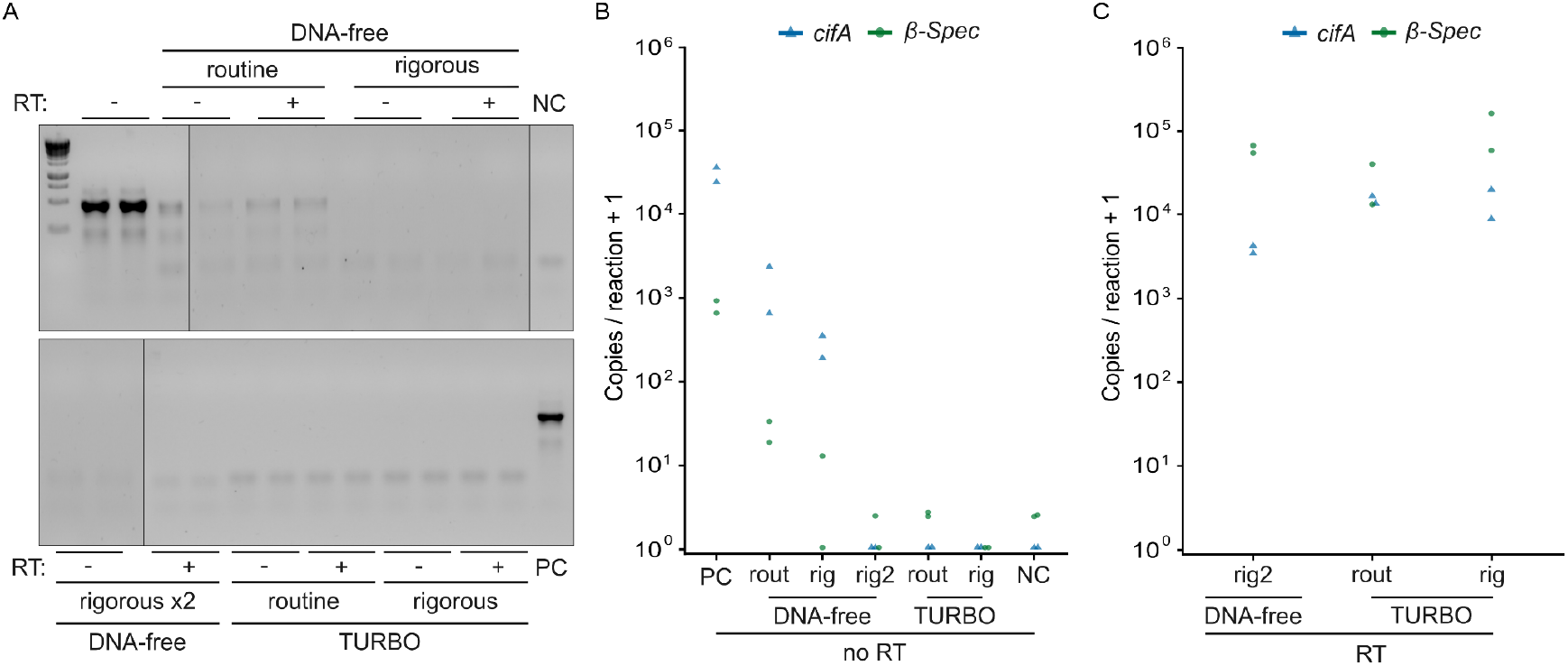
Treatment with TURBO DNase can completely remove contaminating DNA from RNA samples. **(A)** We could not amplify *D. melanogaster* 28s rDNA in RNA treated with either the rigorous DNA-free or any TURBO DNase protocols. In contrast, the DNA-free routine protocol showed a visible 28s-associated amplicon. Black vertical lines indicate locations where the gel was cropped for display. **(B)** All DNase treatment protocols reduced *cifA* and *β-Spec* gDNA copy numbers compared to non-DNase-treated controls. We detected no *cifA* gDNA in the DNA-free rigorous x2, TURBO routine, and TURBO rigorous protocols. Only the TURBO rigorous protocol completely removed *β-Spec* gDNA. **(C)** Among the protocols that achieved complete *cifA* gDNA removal, *cifA* and *β-Spec* RNA concentrations after reverse transcription are similar. Raw gel images and data are available in **Data S10** and **Data S11**.

## Discussion

CI strength directly influences *Wolbachia* prevalence in natural and applied insect populations (Hoffmann et al., 1990, 2011), and stronger CI often correlates with higher *cifB* transcript levels (Shropshire et al., 2022, 2021a). The naturally low abundance of *cifB* transcripts (Lindsey et al., 2018), however, necessitates the pooling of tissues from 15 or more individuals to achieve reliable quantification through standard RT-qPCR procedures. Here, we introduce four RT-ddPCR assays that overcome this obstacle. Below, we discuss the appropriate application for each assay, strategies for further improving assay sensitivity and precision, and generalizability beyond *w*Mel in *D. melanogaster*.

To measure *cif*-gene expression, we developed two distinct assays for each *cif* gene, *cif*/spike and *cif*/*β-Spec*, which enable different normalization strategies. The *cif*/spike assays control for technical variability by introducing a synthetic spike-in RNA to each sample prior to RNA extraction. We assume that any loss of this spike-in RNA during downstream processing (e.g., purification, DNase treatment, cDNA synthesis) is proportional to the loss of endogenous *cif* transcripts. By measuring the recovery of the spike-in RNA via RT-ddPCR, we calculate processing efficiency for each sample. Multiplying raw *cif* counts against processing efficiency corrects for technical variation introduced during sample handling. In contrast, the *cif*/*β-Spec* assay normalizes *cif*-transcript counts to those of a stably expressed *D. melanogaster* housekeeping gene, *β-Spec* (Hu et al., 2017; Shropshire et al., 2021a). This method controls for biological variation, such as the amount of starting tissue or overall host transcriptional activity. This relative quantification is particularly useful for assessing the density of *cif* transcripts relative to host-transcript levels. Before using this assay, it is crucial to confirm that *β-Spec* expression remains stable across all experimental conditions and treatment groups. If *β-Spec* levels are variable, an alternative housekeeping gene should be selected.

We defined the limit of detection for each assay as the upper limit of the 95% confidence interval of the no-template controls (rounded up); this value establishes the threshold below which a signal is indistinguishable from background noise. We calculated limits of detection as 1 *cifA* copy for both assays, 3 *cifB* copies for the *cifB*/spike assay and 1 *cifB* copy for the *cifB*/*β-Spec* assay. Since these values are lower than the transcript quantities measured in dilutions equivalent to individual testes pairs, we conclude these assays are sufficiently sensitive to measure even rare *cif* transcripts from individual insect tissues. However, the limit of detection can vary between experiments depending on background signal levels, which are influenced by lab cleanliness and handling techniques. Therefore, we strongly recommend including both no-template and gDNA elimination controls in every run. Furthermore, while both assays are accurate and precise, precision is marginally lower when counting rare targets. This result is consistent with our prior measurements of *Wolbachia* abundance with ddPCR (Njogu et al., 2025), and is an expected consequence of stochastic variation during pipetting with rare targets and low volumes. Although the assays are highly sensitive and precise, their performance can be further improved by increasing the number of molecules analyzed. This can be achieved by eluting the RNA in a smaller volume than the current 25 μL, increasing the current 2 μL of cDNA per reaction to the maximum allowable volume of 7.5 μL for the *cif*/spike or 8 μL for the *cif*/*β-Spec* assays, and increasing the number of droplets analyzed by performing multiple RT-ddPCR reactions for each sample.

While the *cif*/*β-Spec* RT-ddPCR assays are specific to *D. melanogaster*, the *cif*/spike assays do not measure any host transcripts, making them host-independent. Therefore, we expect the *cif*/spike assays to be applicable to biocontrol programs that deploy *w*Mel *Wolbachia* in *Ae. aegypti* mosquitoes to control viral pathogens such as dengue and Zika (e.g., Utarini et al., 2021). Since strong CI facilitates *w*Mel’s spread, and weak CI can lead to failed public health initiatives (e.g., Ross et al., 2019), these assays may enable efforts to monitor *cif*-transcript levels in field-caught mosquitoes as a proxy for CI-strength variation. Furthermore, we designed the *cif* oligos with homology to at least 39 *cifA* variants (from 32 *Wolbachia* strains) and 34 *cifB* variants (from 27 *Wolbachia* strains). This extends application of these assays to an array of *Wolbachia*-host systems, including agricultural pests like *Delia radicum* (Lopez et al., 2018; Sontowski et al., 2022), *Liriomyza huidobrensis* (Hidayanti et al., 2022; Ohata et al., 2025), and *Rhagoletis cingulata* (Schuler et al., 2016) that destroy crops such as cabbage, peas, beans, and cherries; the livestock pest *Haematobia irritans* (Madhav et al., 2020); and *Hofmannophila pseudospretella*, which damages stored cereals, fabrics, and dried fruit. We recommend performing pilot experiments to confirm the assays’ suitability before applying it to any new system of interest.

## Materials & Methods

### Insect lines, care, and maintenance

We performed experiments using *w*Mel-bearing *D. melanogaster* from the y^1^w^*^ stock (BDSC 1495). We maintained flies under a 12:12 light:dark cycle at 23°C within a *Drosophila* incubator (Percival DR-36VL) using standard narrow *Drosophila* vials (Flystuff 32-113RL) containing 7 mL to 10 mL of fly food (Detailed protocol in Wheeler et al., 2024). We anesthetized adult flies with CO_2_ during experiments. We periodically tested *Wolbachia* cytotypes by extracting DNA from pools of three randomly sampled flies from each stock using a SquishBuffer method. This was followed by PCR amplification of the *Wolbachia* surface protein gene and the host 28s rDNA gene and gel electrophoresis (Detailed protocol in Cooper and Shropshire, 2024).

### Sample collection, RNA extraction, and RNA processing

We detail the following methods in full on protocols.io (Van Vlaenderen & Shropshire 2025a). To collect samples, we anesthetized non-virgin males with CO_2_, dissected their testes in 1x RNase-free PBS (Fisher Bioreagents, BP3994), and transferred tissue to 2 mL centrifuge tubes (Eppendorf, 05414203) containing 800 µL of chilled TRIzol (Invitrogen, 15596026) and three 2.8 mm ceramic homogenizing beads (VWR, 10158-554). We immediately homogenized samples in a bead-mill homogenizer at 1,500 rpm for 2 minutes (Benchmark Scientific, BeadBlaster 96), centrifuged them to bring the contents to the bottom of the tube, and froze the samples at −80°C until processing.

We extracted RNA using TRIzol:Chloroform phase separation. We initially thawed samples, homogenized them again for 2 minutes at 1,500 rpm in a bead mill, and incubated them for 5 minutes at room temperature. We diluted a synthetic spike-in RNA sequence (TATAA Biocenter, RS25SI) 1:64 and added 1 µL to each sample. After adding 160 µL of Chloroform (Thermo Fisher Scientific, 032614.K2), we incubated samples for 5 minutes at room temperature, centrifuged them at 4°C for 15 minutes at 12,000 × *g*, and collected the upper aqueous phase. We added 3 µL of glycogen (20 µg/µL; Invitrogen, 10814010) and 400 µL of isopropanol (Thermo Fisher Scientific, 327272500) to each sample, incubated them for 10 minutes at room temperature, and centrifuged them at 4°C for 20 minutes at 12,000 × *g*. We then discarded the supernatant and washed the pellet four times with 500 µL of 75% ethanol (Koptec, V1001), performing intermittent 5-minute centrifugations at 4°C and 7,500 × *g*. After air-drying the RNA pellet, we dissolved the RNA in 25 µL of low EDTA TE buffer (Quality Biological, 351-324-721). We performed reprecipitation to further clean the sample by adding 62.5 µL of 200 proof ethanol and 2.5 µL of 3 M NaAc (pH 5.2) to each sample, and stored them overnight at −20°C. Following a 30-minute centrifugation at 4°C and 21,000 × *g*, we washed the pellets three times with ice-cold 75% ethanol, as described above. After air-drying the RNA pellet, we dissolved the RNA in 25 µL of low EDTA TE. We used a NanoDrop 1C Spectrophotometer (Thermo Fisher Scientific, ND-ONE-W) to evaluate the quality of each sample and a Qubit 4 Fluorometer (Thermo Fisher Scientific, Q33238) with RNA High Sensitivity (HS) Assay Kit (Invitrogen, Q32855) to measure RNA quantity.

We performed DNase treatment using three protocols from the DNA-free kit (Invitrogen, AM1906) and two protocols from the TURBO DNase kit (Invitrogen, AM1907). We followed the manufacturer’s recommendations for each protocol. For the DNA-free kit, we tried the “routine” and “rigorous” protocols, and we also performed the rigorous protocol twice. For the TURBO DNase kit, we performed both the “routine” and “rigorous” protocols. For both kits, the routine protocol involved adding 1 µL of DNase and 2 µL DNase buffer to 20 µL of RNA, incubating at 37°C for 30 minutes, adding an inactivation reagent, and transferring the supernatant after pelleting the inactivation reagent. The rigorous protocol differed only in that we added another 1 µL of DNase after the initial 30-minute incubation and incubated for an additional 30 minutes. We initially determined if DNA was removed from the reaction by performing PCR for the 28s rDNA of *D. melanogaster* and gel electrophoresis (Detailed protocol in Cooper and Shropshire, 2024). First-strand cDNA synthesis was performed using the SuperScript IV VILO Master Mix (Invitrogen, 11756050). cDNA was either immediately used for RT-ddPCR or stored at −80°C.

### Oligo design for ddPCR

We designed three new RT-ddPCR oligo sets (primers and probes) for this study, targeting *cifA, cifB*, and *β-Spec* (**Table 1**). To design the *cif* oligo sets, we retrieved relevant sequences from an in-house collection of *Wolbachia* genomes using BLAST (e-value 1e-10) with *w*Mel *cif* sequences as queries. We generated a multiple sequence alignment (MSA) using Muscle5 (Edgar, 2022) and obtained a consensus sequence with variable sites represented as Ns using Geneious Prime v2024.0.5. We used Primer3web v4.1.0 (Koressaar et al., 2018) to design oligos based on the consensus sequence. We iteratively refined the sequence list through multiple rounds of MSA and oligo design until we acquired oligos matching the criteria below. We used Primer-BLAST to determine the number of *cif* variants and *Wolbachia* strains that the *cif* oligos are homologous to in the NCBI nr database. We designed the *β-Spec* oligo set using Primer3web based solely on the *D. melanogaster* sequence (GenBank M92288).

**Table 1.**
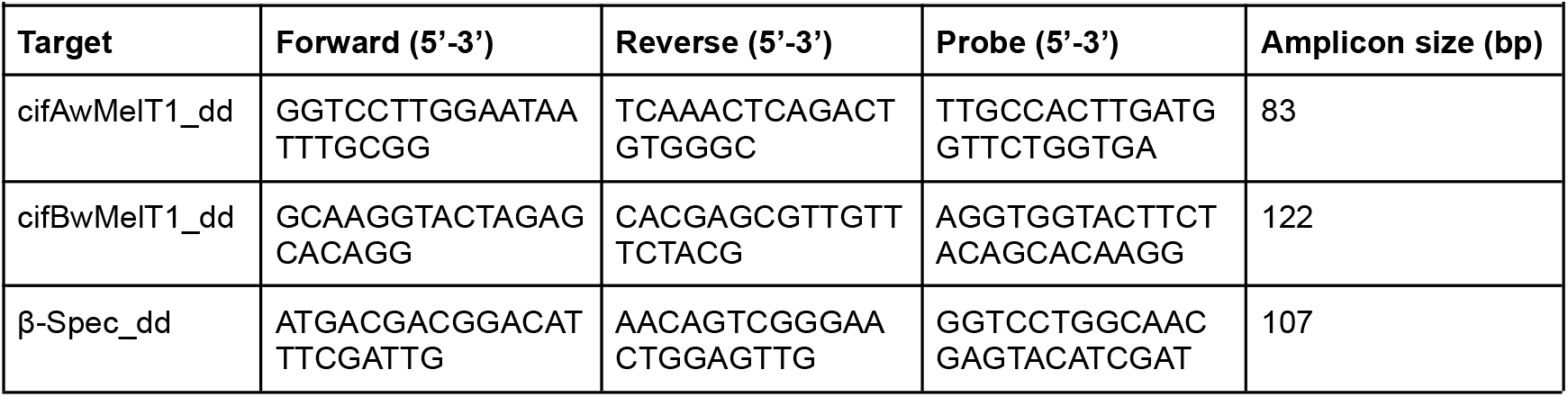
RT-ddPCR oligos for measuring *cifA* and *cifB* mRNA.

Primers were between 18 and 22 base pairs (bp) long, and yielded amplicons ranging from 70 to 150 bp. They also had a GC content of 40 to 60%, a GC clamp of 2, a melting temperature (Tm) of 58 to 62°C, a maximum Tm difference of 2°C, and a maximum of 3 consecutive identical nucleotides (poly-X). Probes were 18 to 30 bp in length with a Tm of 65 to 70°C, a GC content of 30 to 80%, and a maximum of 5 consecutive identical nucleotides. We performed Tm calculations using the SantaLucia algorithm (SantaLucia, 1998) with specified ion and dNTP concentrations. To ensure specificity, we analyzed primers against human (taxid: 9606), *D. melanogaster* (taxid: 7227), and *Wolbachia* (taxid: 953) genomes using Primer-BLAST. We purchased premixed primer-probe sets from Bio-Rad at a 900 nM:250 nM ratio. We purchased proprietary oligos for the synthetic spike-in RNA from TATAA Biocenter (RS25SI).

### RT-ddPCR

We conducted all RT-ddPCR reactions in 20 µL volumes. We prepared *cifA*/spike and *cifB*/spike duplex reactions using 10 µL of ddPCR Supermix for Probes (no dUTP) (Bio-Rad, 1863023), 1 µL of the relevant *cif* oligo set, 1 µL spike-in RNA primer mixture, 0.5 µL spike-in RNA probe, 5.5 µL of nuclease-free water, and 2 µL of cDNA template. We prepared *cifA*/*β-Spec*, and *cifB*/*β-Spec* duplex reactions using 10 µL of ddPCR Supermix for Probes (no dUTP) (Bio-Rad, 1863023), 1 µL of each relevant oligo set, 6 µL of nuclease-free water, and 2 µL of cDNA template. After preparation, we sealed PCR plates with an adhesive film (Bio-Rad, MSB1001), vortexed them for 10 seconds to mix (Four E’s Scientific, MI0101002), and centrifuged them for 2 minutes at 2,204 × *g* (Eppendorf, 5430-R). We then removed the adhesive film and transferred 19.5 µL of each reaction mixture to a droplet generation cartridge (Bio-Rad, 1864007). We added 70 µL of droplet generation oil for probes (Bio-Rad, 1863005) to the adjacent well, placed a gasket (Bio-Rad, 1864007), and positioned the cartridge in a droplet generator (Bio-Rad, QX200) to create droplets. Next, we transferred the 40 µL droplet wells to a 96-well plate, sealed them with a heat-activated adhesive film (Bio-Rad, 1814040 and PX1), and placed them in a thermal cycler (Bio-Rad, C1000 or S1000) for PCR. Reaction conditions varied across experiments, as described in the results section. After PCR, we maintained samples at 12°C until analysis on a droplet reader (Bio-Rad, QX200). We excluded samples that yielded fewer than 10,000 droplets. We present a detailed RT-ddPCR protocol for *cif*-transcript analysis on protocols.io (Van Vlaenderen & Shropshire 2025).

### Statistical analysis and figure generation

We used QX Manager software (v.2.1.0; Bio-Rad) to calculate RT-ddPCR target concentration and confidence intervals, and to produce RT-ddPCR plots. We performed all other statistical analyses in R (v.4.4.1) using RStudio (v2024.04.2). To create plots, we used the “ggplot2” package (v.3.4.4) in R (R Core Team, 2024; Wickham, 2016). Finally, we used Inkscape (v.1.3.2; Inkscape Developers) to modify figure aesthetics.

## Acknowledgments

We gratefully acknowledge Helene Hartman, Lara Parada-Tixe, and Abby Klebe for their support in the laboratory. Constructive feedback from Alphaxand Njogu, Helene Hartman, and other members of the Shropshire Lab greatly enhanced the quality of this work. We also extend our appreciation to the staff in Lehigh University’s Department of Biological Sciences for their valuable support.

## Contributions (CRediT Classification)

<u>Conceptualization</u>: JDS. <u>Data curation:</u> LVV, WRC, JDS. <u>Formal analysis</u>: LVV, WRC, JDS. <u>Funding acquisition</u>: JDS. <u>Investigation</u>: LVV. <u>Methodology</u>: LVV, JDS. <u>Project administration</u>: JDS. <u>Resources</u>: JDS. <u>Supervision</u>: JDS. <u>Visualization</u>: LVV, JDS. <u>Writing – Original draft</u>: LVV, JDS. <u>Writing – Review & editing</u>: LVV, WRC, JDS.

## Data availability

All data are publicly available in the supplement of this manuscript.

## Supporting Information

**Data S1. RNA extraction and purification measurements**.

**Data S2. FASTA file containing *cifA* sequences homologous to the *cifA* RT-ddPCR oligos**.

**Data S3. FASTA file containing *cifB* sequences homologous to the *cifB* RT-ddPCR oligos**.

**Data S4. *cifA*/spike RT-ddPCR annealing temperature measurements**.

**Data S5. *cifB*/spike RT-ddPCR annealing temperature measurements**.

**Data S6. *cifA*/spike RT-ddPCR dilution series measurements**.

**Data S7. *cifB*/*spike* RT-ddPCR dilution series measurements**.

**Data S8. *cifA*/*β-Spec* RT-ddPCR dilution series measurements**.

**Data S9. *cifB*/*β-Spec* RT-ddPCR dilution series measurements**.

**Data S10. DNase treatment raw gel images**.

**Data S11. DNase treatment RT-ddPCR results**.

