## Supplementary figures and images for "Counting cytoplasmic incompatibility factor mRNA using digital droplet PCR"

### Data-S10.png

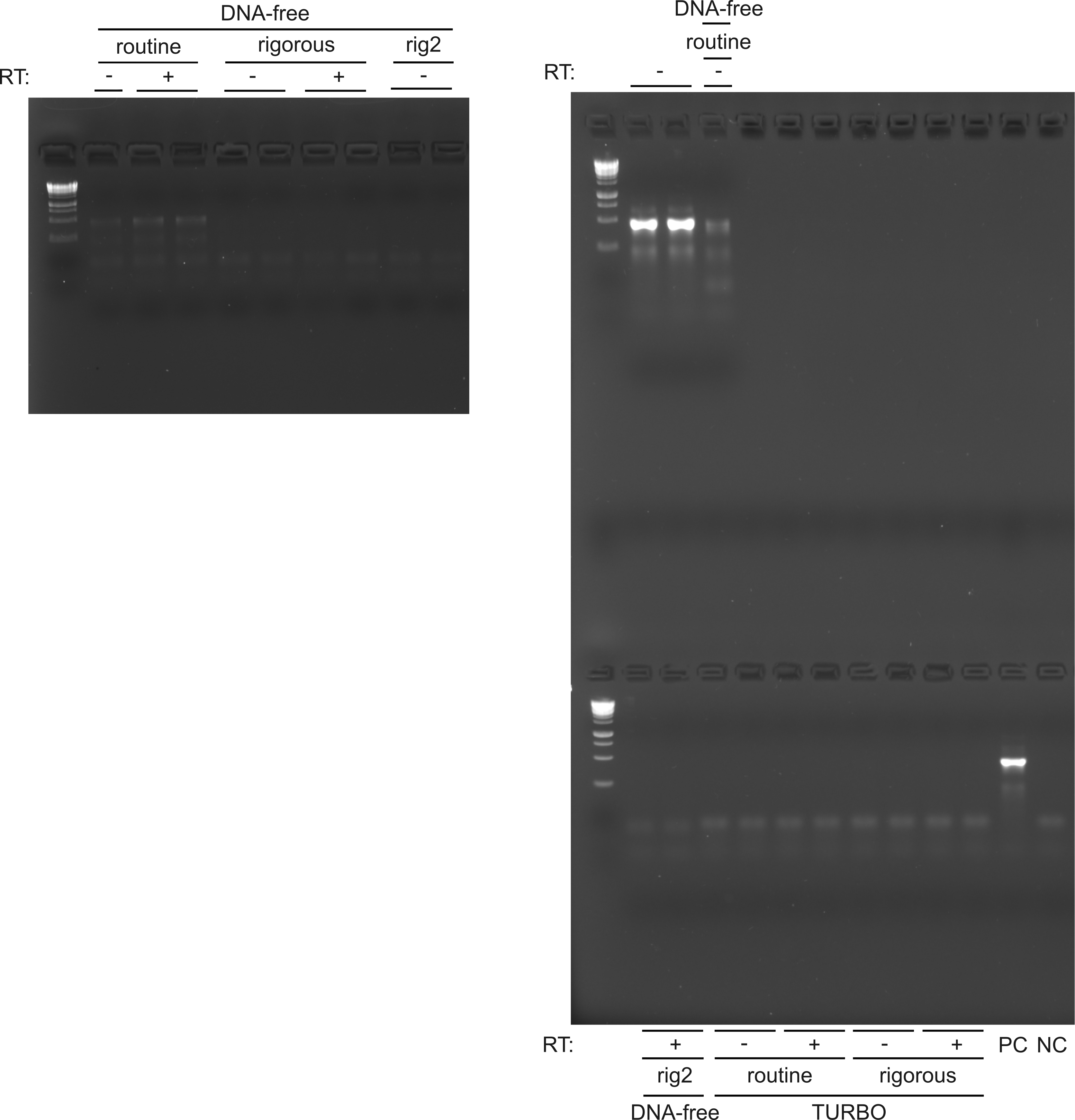
